# Quantifying B-cell Clonal Diversity In Repertoire Data

**DOI:** 10.1101/2022.12.12.520133

**Authors:** Aurelien Pelissier, Siyuan Luo, Maria Stratigopoulou, Jeroen EJ Guikema, Maria Rodriguez Martinez

**Author notes:** Equal contribution (Aurelien Pelissier, Siyuan Luo).

## Abstract

The adaptive immune system has the extraordinary ability to produce a broad range of immunoglobulins that can bind a wide variety of antigens. During adaptive immune responses, activated B cells duplicate and undergo somatic hypermutation in their B-cell receptor (BCR) genes, resulting in clonal families of diversified B-cells that can be related back to a common ancestor. Advances in high-throughput sequencing technologies have enabled the high-throughput characterization of B-cell repertoires, however, the accurate identification of clonally related BCR sequences remains a major challenge. In this study, we compare three different clone identification methods on both simulated and experimental data, and investigate their impact on the characterization of B-cell diversity. We find that different methods may lead to different clonal definitions, which in turn can affect the quantification of clonal diversity in repertoire data. Interestingly, we find the Shannon entropy to be overall the most robust diversity index in regard to different clonal identification. Our analysis also suggests that the traditional germline gene alignment-based method for clonal identification remains the most accurate when the complete information about the sequence is known, but that alignment-free methods may be preferred for shorter read length. We make our implementation freely available as a Python library `cdiversity`.

## Introduction

Antibodies are protective proteins produced by B cells in response to the presence of foreign pathogens, they have an exceptional ability to recognize a wide variety of target antigens and can display exquisite binding specificity [1]. To ensure broad antigen recognition, they undergo several rounds of maturation where they gain increased affinity, avidity, and anti-pathogen activity. Briefly, B cell receptors (BCRs) are assembled through the rearrangement of the V, D, and J gene segments, coupled with stochastic insertions and deletions of nucleotides at the gene boundaries, i.e. at the V and D, and the D and J gene *junctions* [2]. The junctions between the V, D, and J gene segments are known as the complementary determining region 3 (CDR3). This region is the most diverse part of the BCR sequence and plays a crucial role in determining the binding specificity to foreign antigens.

Once a BCR has been formed, B cells are exposed to antigens in the secondary lymphoid organs and undergo affinity maturation in microanatomical structures known as Germinal Centers (GCs). Through antigen-driven competition, selected B cells receive secondary signals that direct them to undergo further rounds of cellular replication and BCR diversification through somatic hypermutation (SHM). Through this process, B cells with higher affinity to the target antigen are preferentially selected to further expand, while the ones with lower affinity undergo programmed apoptosis. This process results in the progressive expansion and evolution of the initial pool of founder cells into distinct groups of clonally related B cells (referred to as B-cell clones) that compete against each other for antigen-mediated survival signals. Through an accelerated Darwinian process of diversification and selection, some of these clones expand significantly and can become dominant, while others disappear [3].

Because of the stochastic nature underlying clonal selection coupled with the randomness associated with experimental BCR sampling and sequencing, it is common to observe a fraction of B cells without any clonally related B-cell in a repertoire. In this manuscript, we refer to this group of B cells as *singletons,* and use the term *non-singletons* to refer to B cells that have other clonally related B cells. Of course, the distinction between *singletons* and *non-singletons* depends on a user-defined threshold to separate clonally related and unrelated cells, as will be discussed in Section 1.2.

The constant evolution and antigen-driven selection of B-cell repertoires make the sequencing and analysis of BCRs a valuable tool to identify the immune status of an individual. Indeed, as memory B cells produced during short-lived immune episodes can survive for a very long time, sometimes for the entire lifetime of a person, their analysis also reveals information about the past and current pathogens encountered by an individual [4, 5]. Furthermore, beyond infections, the analysis of B-cell repertoires can provide valuable fingerprints of an individual’s immunological status, and enable the diagnosis of complex diseases, chronic inflammatory conditions, allergies, response to vaccination, etc [6, 7, 8, 9, 10, 11, 12, 13]. Advances in Adaptive Immune Receptor Repertoire Sequencing (AIRR-Seq) technologies have considerably increased the amount of repertoire data that is available for analysis and improved our understanding of the dynamics of B-cell repertoires in both individuals and populations.

The analysis of B-cell repertoire typically starts by grouping BCR sequences into clones of related B cells. The reconstruction of phylogenetic lineages of clonally related B cells provides information about the evolutionary paths that led to the development of functional antibodies and has been shown to be useful to understand the progression of diseases such as chronic infections, autoimmune diseases or cancer. Still, this previous work focused on diseases that revolve around a single stereotyped mutation. In most cases, immune repertoire data are distinct across individuals in humans and mice with respect to their clonal composition [14], and even between identical twins [15]. The CDR3 sequences from both unimmunized and immunized or vaccinated individuals only show small sequence overlap and oftentimes no sequence overlap at all [16, 17, 18]. This makes a direct sequence repertoire comparison across individuals inadequate to identify robust immune-repertoire-based signatures.

A more promising approach to comparing immune repertoires across individuals focuses on the investigation of sequence-independent quantifiers such as clonal diversity indices. These quantifiers offer the possibility of correlating immune repertoire diversity to immunological status and, in doing so, readily allow for immune-repertoire-based comparisons across individuals. Nevertheless, there is still substantial ambiguity and a lack of quantitative understanding about the effectiveness of the diversity metrics to reliably capture status-specific information from immune repertoires. Realistic measures of diversity should reflect not only the relative abundances of clones, but also the main differences between them [19, 20]. Furthermore, the use of specific diversity indices, such as the Shannon [21] or Simpson [22] diversity indexes, may yield qualitatively different results in different contexts [19, 20]. Intuitively, that is because these indices do not put the same weights on the clone abundances in the repertoire. For example, the richness is most sensitive to the rarest clones, while Simpson index (probability that two randomly selected individuals belong to different species) or dominance are affected mainly by the most common clones.

The limitations associated with individual metrics have supported the practice of aggregating multiple indices for immunological classification. For instance, the Hill-based diversity profiles are composed of a continuum of single diversity indices and can facilitate a global quantification of the immunological information contained in immune repertoires [23, 24]. Still, the estimation of these indices may vary drastically depending on the sequencing depth of the sequencing experiment. For example, species richness quantifies the number of species in a sample, and is particularly affected by the sample size. Several bias estimators have been proposed to correct for incomplete sample information (Chaos estimator [25, 26]). However, all these estimators present vulnerabilities that limit their applicability to immune repertoires with variable sequencing depth [27]. Chaos for example rely heavily on the correct quantification of singletons and doubletons, which is often prone to error in the context of B-cell repertoire.

In fact, another challenge in the analysis of B-cell repertoire data is the grouping of BCR into clones. Theoretically, a group of clonally related B cells represents a group of B cells descending from the same common ancestor (aka founder cell). In a real experiment, the founder cell has long disappeared, being replaced by better-adapted descendants. Therefore, in most cases reconstructing phylogenetic trees amounts to inferring trees where the founder cell is unknown. In practice, B cells with similar characteristics, such as same gene segments and similar CDR3, are grouped together in the same clone. This *empirical definition* of clone poses several challenges, the first being the arbitrariness of the choice of BCR properties used to define a clone, and second, the subjective choice of metric and threshold used to separate clonally related and unrelated B cells. Importantly, the optimal way to group clones may differ depending on the dataset.

In this context, a variety of methods have been proposed to (semi-)automatically identify clones from a set of BCR sequences. Some are based on probabilistic models that infer a hypothetical unmutated common ancestor to be used as tree root, which enables the inference of rooted trees interpreted as clones [28, 29]. The most common techniques rely on CDR3 sequence similarities as well as the alignment of BCRs to reference V and J gene germline sequences [30, 31, 32, 33, 34]. As these alignments are prone to error, some recent approaches leverage natural language processing (NLP) techniques to define similarity indicators independent of these gene alignments [35].

### Our contribution

Each method for clonal identification depends on arbitrary choices of BCR sequence features, sequence-based distances and thresholds, and therefore, may lead to substantially different clonal groupings – besides displaying high variability in computational complexity and robustness. This variability might affect the estimation of clonal diversity which, as we discussed earlier, is crucial for the global analysis of B-cell repertoire data. In this work, we consider different definitions of clones and investigate the consistency and robustness of commonly used diversity indices at different sequencing depths and across different samples and technical replicates. By objectively comparing the performance and consistency of both clone definitions and diversity indices in various experimental contexts, we aim to investigate in which way the different empirical definitions of clones affects diversity analysis and the qualitative biological conclusions extracted from it. Finally, in order to facilitate the use of different clonal identification methods and diversity metrics for future studies, We make our implementation freely available as a python library (`cdiversity`).

## 1 Results

### 1.1 B cell repertoire data

To perform a diversity analysis and characterize the influence of the different metric and definition choices, we collected three different B-cell repertoire datasets from different biological contexts:

- **Simulated data**. We collected artificial repertoires with known clonal relationships [35], generated by randomly adding mutations based on the learned lineage tree topologies from a multiple sclerosis (MS) study [36]. The repertoires thus exhibit a wide range of sequence and junction lengths and diverse clone sizes. While the artificial generation may bias the repertoire towards certain patterns used in its generation, they provide essential information about both the B cells clonal relationships and their lineage history (ground truth information). From this, we can estimate exact diversity indices metrics, information that is inaccessible in experimental datasets.
- **Germinal center data**. The second dataset is a collection of B-cell repertoires from 10 individual GCs within the same lymph node of a patient with chronic sialadentitis [37], with two replicates per GCs. Importantly, the first 70 nucleotides of the V gene segments are missing due to the experimental design (primers were binding within the FR1 region, which was thus removed during the processing of the reads in order to avoid PCR or sequencing errors [38]). This limitation makes this dataset particularly interesting for our study, as it enables us to asses the impact of an uncertain V gene assignment in diversity estimation. In addition, as GCs can be considered semi-independent evolutionary structures with limited cell exchanges, they exhibit a high variability in B-cell diversities even within the same lymph node [30]. Therefore, the comparisons of the technical replicates from the same GC will allow us to establish confidence values to our inferred diversity estimators, as samples extracted from the same GC are expected to exhibit a similar degree of diversity compared to samples from other GCs.
- **Vaccination data**. Our third dataset comes from a study of hepatitis B-associated chronic infection and vaccination responses [6]. This dataset characterizes the different B-cell repertoire landscapes of individuals shortly after vaccination and/or infection compared with controls (non-vaccinated and non-infected individuals). The dataset contains 27 samples from controls, infected as well as pre- and post- (2 weeks) vaccinated individuals.

### 1.2 Different Clonal identification methods yield inconsistent B-cell groups

The first step in the analysis of B-cell repertoire data is the grouping (or clustering) of BCRs into clonal families. In this work, we focus on three approaches to identify clones previously described in the literature.

- **Junction-based methods:** In this method, B cells are assigned to the same clone if and only if their receptors share the same CDR3 sequence. It has the advantage of being computationally simple and of eliminating the ambiguity in defining arbitrary clustering thresholds. On the other hand, since the junction of clonally related B cells typically exhibit small sequence differences due to SHM, this approach will inevitably split branches of the same lineage into different clones, leading to an inflation of the diversity metrics.
- **Alignment-based methods:** A more commonly used approach relies on the junction sequence *and* the V and J assignments [39] (we refer to this method as VJ & Junction). Namely, B cells are assigned to the same clone if their receptors share the same V and J gene segments and exhibit an X% of CDR3 sequence similarity, where the similarity is assessed with the Levenshtein distance [40] and the threshold is set around 90%, with some variability depending on the dataset. These methods allow for a small sequence divergence in clonally related cells due to potential insertions and deletions through the subsequent rounds of B-cell diversification through SHM. In practice, the X% threshold is adjusted for each dataset independently. There are several ways of doing so. A first, intuitive approach consists in computing the distances between pairs of junctions from B cells with the same V and J gene segments (Figure 1A). The distribution of pairwise distances is expected to be the mixture of two distributions, one corresponding to distances between members of the same clone (non-singleton sequences) and the second corresponding to distances between clonally unrelated sequences (singletons). The value that separates the two modes of the distribution can then be used as a threshold to separate both clonally-related and unrelated sequences [41] (Figure 1A). The bi-modality-based threshold has however a high computational cost. An alternative method is based on the assumption that clones do not span multiple individuals. Hence, sequences randomly sampled from multiple unrelated individuals, i.e. *negation sequences,* can be introduced and used to define a threshold by computing the distribution of distances between negation sequences and their closest counterparts within the considered individual [42] (Figure 1A). In practice, a threshold is chosen that allows a fraction of false-positive sequences roughly equal to a tolerance *δ* to be below the chosen threshold. This heuristic aims for high specificity, which is approximately 1 – *δ*. In this work, the threshold was set using a tolerance of *δ* = 1%. Finally, the threshold and the computed CDR3 pairwise Levenshtein distances are used together with a Hierarchical Agglomerative Clustering (HAC) algorithm [43] to further split BCRs with the same V and J gene segments into different clonal groups [39].
- **Alignment free:** As germline gene alignments are error-prone, alignment-based methods might fail to identify clonal relatives accurately, especially when part of the nucleotides are missing in the sequences (as in the GC dataset considered in section 1.3). To overcome this limitation, an alignment-free method that leverages NLP techniques has been recently introduced [35]. Briefly, the method decomposes each BCRs into *k*-mers (substrings of length *k*) and uses the term frequency-inverse document frequency *(tf-idf)* as a weighting scheme that increases proportionally to the number of times a *k*-mer term appears in the document but is offset by the frequency of the term in the corpus. The logic behind this is to emphasize rare and hopefully meaningful terms while reducing the influence of common and uninformative terms. Once a vectorized representation of each BCR has been built, BCR similarities are computed with the cosine distance, which allows for the very fast computation of similarities among text strings. The pairwise distances are then fed into a HAC algorithm to compute the final clusters, with a distance threshold defined from the negation sequences in the same way as with the *alignment-based* method (Figure 1B).

**Figure 1:**
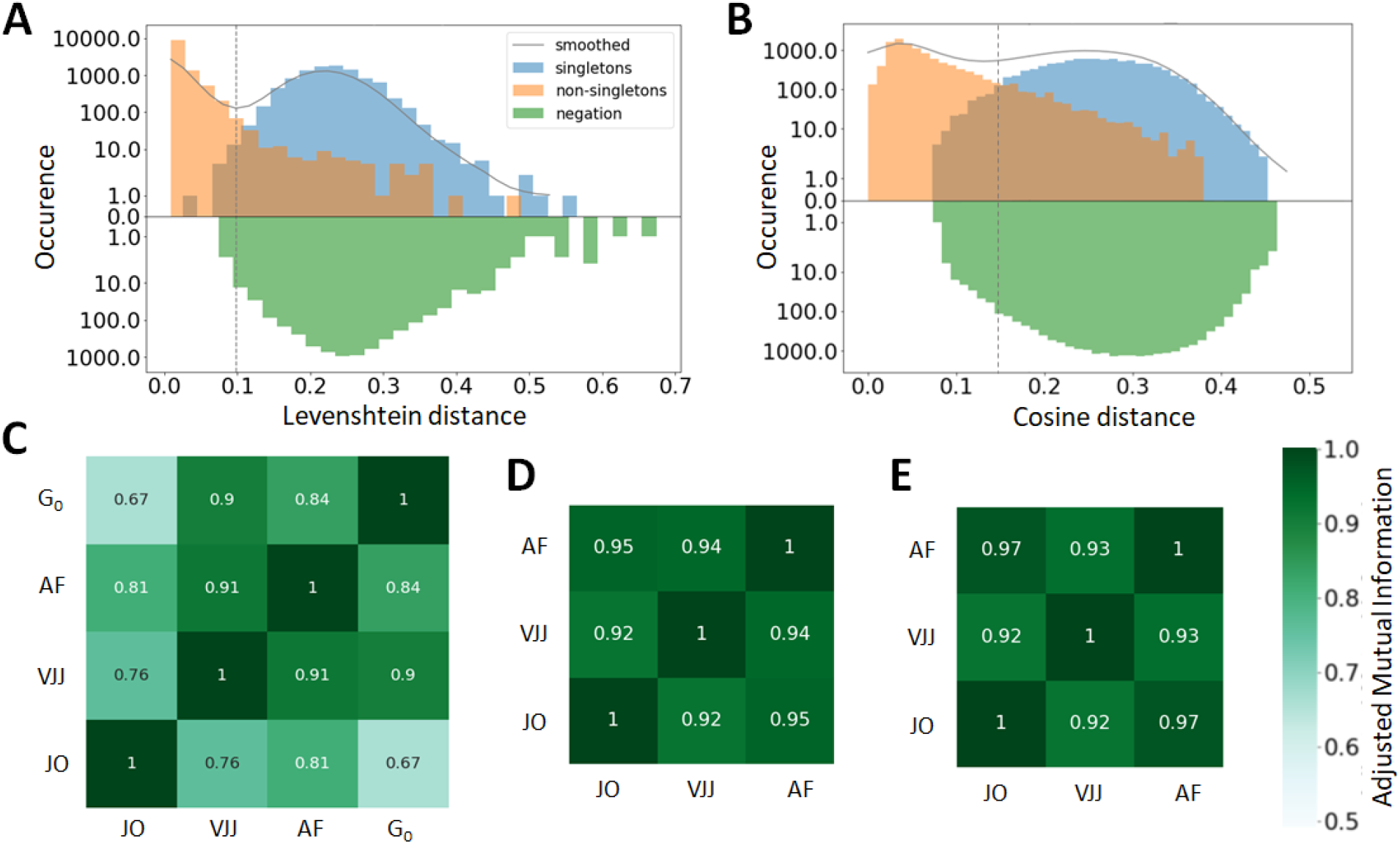
(A&B) Distance to the nearest neighbor sequence distribution, both within the same repertoire (blue and orange) and to the negation sequences (green). The distances of singletons and non-singletons sequences are labelled according to the ground truth, i.e. the clonal groups in the simulated data. Results are shown for both the (A) Alignment-based method, i.e. based on common V-J segments and junction similarity, and (B) alignment-free method, based on the cosine distance computed between vectors representing the frequency of different k-mers on each BCR (*k* = 7). Thus, the metrics used by both methods are not the same. For additional information, the distance threshold used for the clustering by each clonal identification methods is also displayed. (C,D,E) Adjusted mutual information between the clusters obtained from the three methods: *Junction-only* (JO), *VJ & Junction* (VJJ), *Alignment-free* (AF) and *Ground-truth* (G_0_). Results are averaged over all samples and provided for the (C) simulated, (D) germinal center and (E) hepatitis B dataset. The ground truth about clonal clusters is only available in the simulated dataset

A straightforward way to visualize the consistency and performance of the different clonal identification methods is through the classification of singletons. As mentioned earlier, singletons are clonally unrelated sequences, and thus, we expect them to exhibit larger distances to their nearest neighbors in the repertoire than non-singletons sequences. We first run a comparative analysis in the simulated dataset, as we have ground truth information about clonal relationships. Figure 1A&B displays the distance to the nearest neighbors of each B cells in the simulated data, for both singletons (blue) and non-singletons sequences (orange). We observe that both methods failed to identify accurately some of the singletons in the simulated dataset (8% for alignment-free, and 1% for VJ & Junction). These inaccuracies can be visualized with the (i) blue sequences to the left of the vertical line and (ii) orange sequences to the right of the vertical line (note the log-scale on the *y*-axis). Overall, our analysis suggests that the VJ & Junction method (alignment based) is the best at classifying singletons in the simulated data.

Regarding other datasets, while we don’t have information about the true clones, we observed that the methods also disagree within their respective classification of singletons (SI section 1). To better quantify the similarity of the inferred clonal group across methods, we computed the adjusted mutual information (AMI) [44] between the clusters inferred with each method and for each dataset (Figure 1C,D&E). AMI is a variation of mutual information that compares the different partitions produced by different clustering schemes. Furthermore, AMI also corrects for the effect of the agreement between clusters solely due to chance. A value close to 0 indicates no overlap, while a value of 1 corresponds to identical clusters. As expected from the differences in clonal definitions used by the three methods, we observe differences in the clusters inferred by each method. Focusing first on the synthetic dataset for which the ground truth is known (Figure 1C), the VJ & Junction method performs best and achieves an AMI with the ground truth of 0.9, while the junction-only method gives the lowest performance with an AMI of 0.67. Importantly, additional analysis revealed that these differences are not due to chance or subsampling, as the three methods were found to be consistent even across different sample sizes and shuffling (Figure S2 in SI section 2). In fact, the clustering relationship between two given sequences may change when the HAC algorithm and *tf-idf k*-mer representations are being computed on a different repertoire. This supports our hypothesis that diversity quantification significantly depends on the method used for clonal identification.

Interestingly, the AMI between the three methods is higher in the case of the GC and Hepatitis B dataset. (AMI >0.92). That is because in these dataset, there are a few abundant clones with many identical junctions, thus inflating slightly the AMI between the three methods (dominance ~ 10%*vs ~1%* for the simulated dataset).

### 1.3 V and J gene segments may be misaligned, impacting the clonal identification accuracy in alignment-based methods

As mentioned earlier, current conventions for clonal identification methods rely on the correct alignment of the germline V and J gene segments to the BCR sequences. Unfortunately, V gene assignments can be ambiguous, especially for shorter read lengths [45]. More concretely, in the GC dataset, the first 70 nucleotides of V gene segments are missing due to experimental constraints, and therefore, it is possible that a non-negligible portion of V genes is incorrectly called biasing the clonal characterization with the VJ & Junction (alignment-based) method on this dataset. Such limitations, originating from the use of FR1 primers, are common in next generation sequencing experiments [37, 38]. To further investigate this hypothesis, we looked for sequences with ambiguous V and J gene annotations in the IgBlast output [46] (multiple gene matches with equivalent alignment score), and found that 13% of assigned V genes were not unique, as compared to only 0.05% for J genes (Figure 2A).

**Figure 2:**
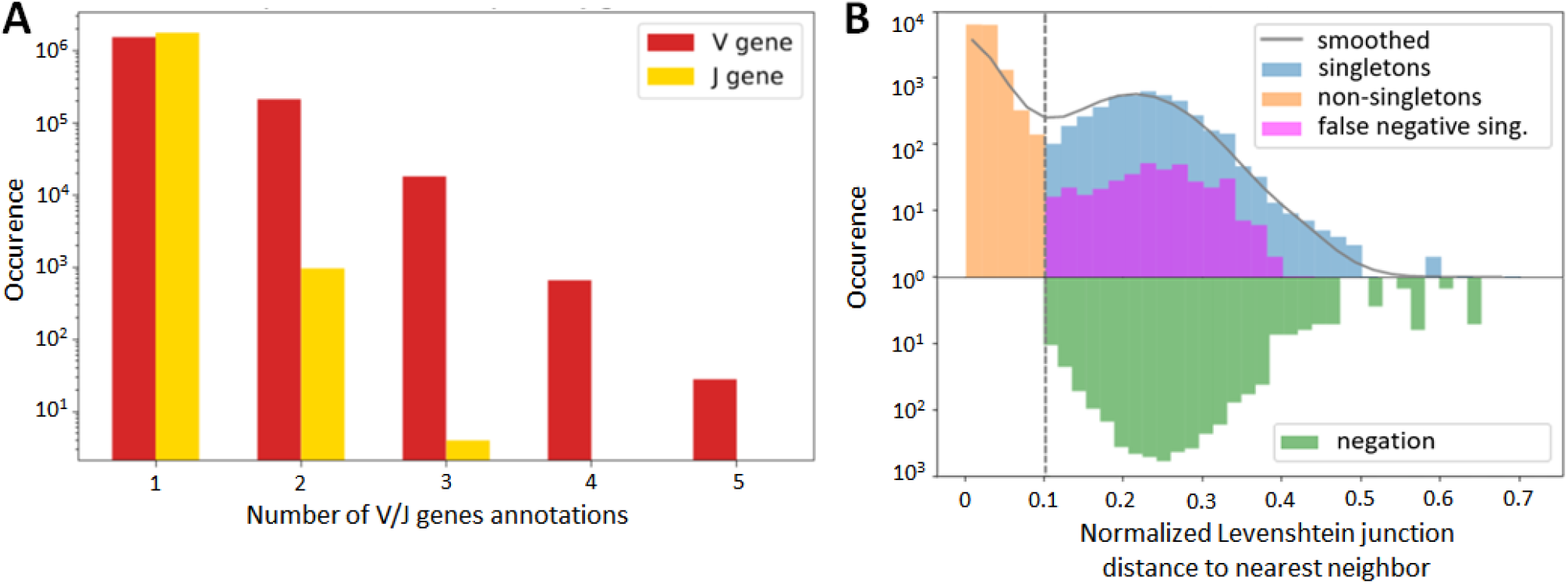
Germline gene alignment and clone identification in the germinal center dataset. (A) B-cell sequence count with more than one match for their V and J gene segment annotations in IgBlast query (equivalent alignment score). (B) Distance to nearest distribution for B-cell sequences in the alignment-based (VJ & Junction method) clonal identification method. False negative singletons originating from V or J gene misalignment are depicted in purple.

To further test for the impact of these potential V gene misalignment, we considered the singleton sequences in the GC dataset. For each singleton identified in our repertoire, we looked at the top 6 V and J gene annotations, and checked how often one of these annotations would lead to an assignment to an existing cluster. We refer to these singletons as potential false negatives, because the method would have *falsely* assigned them as singletons, while it is likely that they belong to an existing clone. Strikingly, we found that 8.8% of singletons inferred by the VJ & Junction method in the GC dataset were potential false negative (depicted in purple on Figure 2B, note the log-scale in the *y* axis). Interestingly, the alignment-free correctly classified 83% of the 8.8% identified false negatives (depicted in purple in Figure S2). This suggests that, while the VJ & Junction method might be optimal when the full sequence information is available, the alignment-free method could be preferable when the germline V gene alignment is ambiguous.

### 1.4 Sensitivity of diversity indices to clonal identification methods

In the previous section, we showed how inferring clonal relationships in B-cell repertoires might markedly vary depending on the method used. We now ask how these inconsistencies affect repertoire comparisons when characterized by different diversity indices. Namely, we calculated sample diversity using various metrics for all samples and all three clonal identification methods. As diversity metrics, we considered dominance [47], richness [48], Simpson index [22], Shannon entropy [21], Hill’s diversity [23] profiles, and some Chao statistical estimators[25, 26, 49] of these accounting for incomplete sample information (see Methods Section 2.4.3).

Generally, we found that although the 3 clone identification methods result in significantly different diversity indices, these indices show similar patterns of variation across samples. Figure S3A shows the Shannon entropy profiles across different samples when clones are computed with the 3 different clonal identification methods. On Figure 3A we grouped replicates together so that we can assess variability across both samples and replicates. These figures clearly illustrate that, although the Shannon entropy values are numerically different depending on the method used to identify the clones, the entropy rank across samples is consistent, and in the case of the GC dataset, the variability across GCs seems higher than the variability between replicates. This implies that some underlying patterns of the repertoire clonal structure are captured by all clone identification methods, and can thus be reflected in the diversity indices. We quantified this trend in terms of the Spearman correlation coefficient of the diversity indices, a Spearman value close to 1 indicates that the two tested methods lead to highly similar diversity-based sample ranks. To quantify this rank similarity in the context of different biological context, we computed the correlation for both cross-method and cross-GC replicate comparisons, for all diversity indices. We reproduce the cross-method analysis on the other two datasets and provide all the computed correlation in Supplementary Table S1.

**Figure 3:**
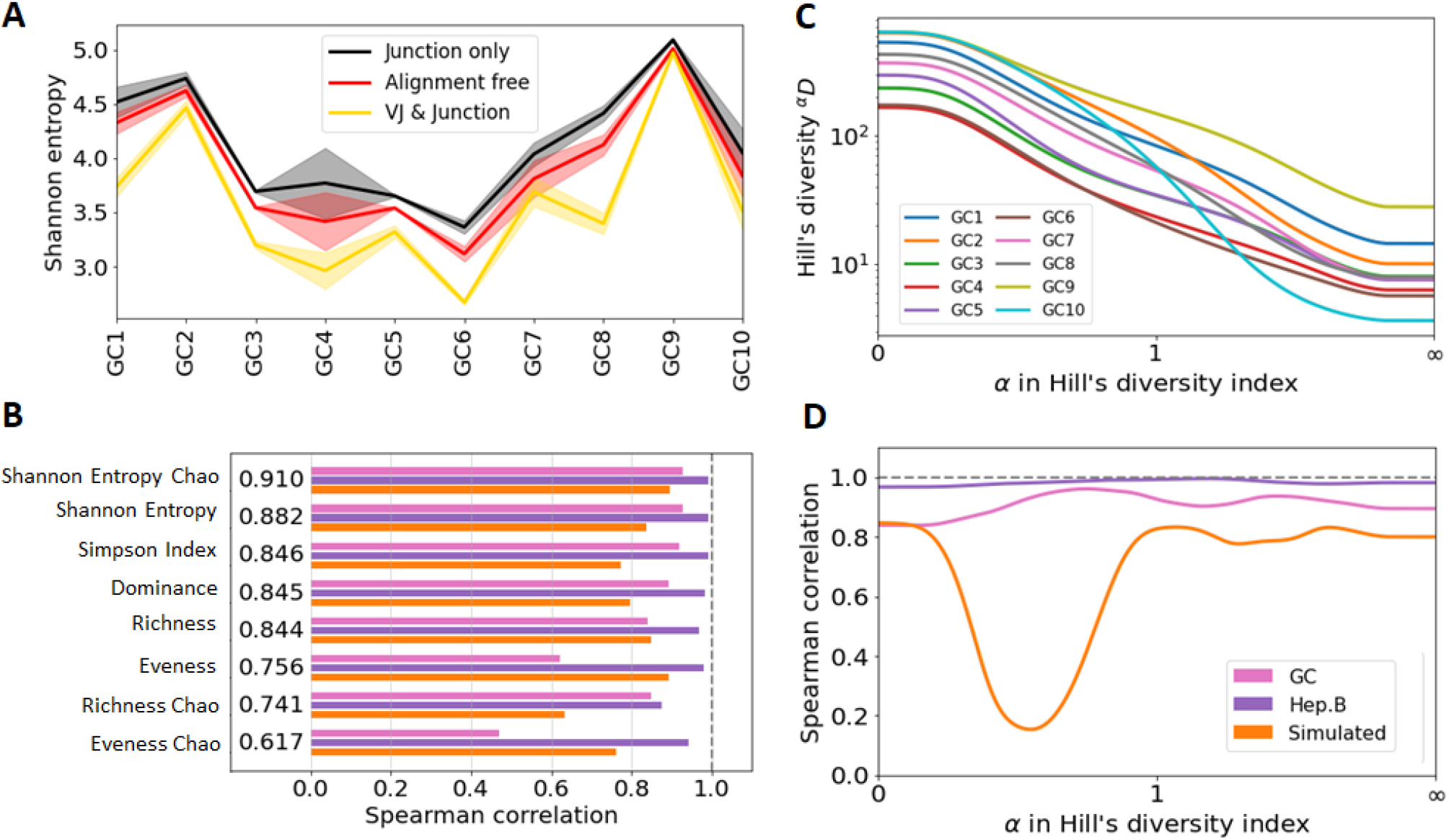
Consistency analysis of diversity indices across different clone identification methods. (A) Shannon entropy diversity index for each GC. As there is two replicates for each GC, the solid line represent the mean value between the two replicates, and the shaded area the min and max value. (B) Spearman correlation between the diversity indices obtained with the three clonal identification methods (mean over pairwise correlation). The computed correlation is shown for the three dataset analyzed in our study and the displayed number is the average correlation across the three datasets. (C) Hill’s Diversity profiles of each sample in GC data with clones obtained from the alignment free method. Note that the *x* axis was transformed by a exponential tangent function for visual clarity. (D) Mean Spearman correlation between the diversity indices obtained with the three clonal identification methods for different values of *α* in the Hill’s diversity frameworks, on the three datasets.

Interestingly, the correlation scores vary drastically depending on the dataset and the task we evaluated. A difference partly explained by the differences in variability of the diversity indices across samples (mean/std ranging from 1 to 80, SI section 5). Intuitively, it is harder to rank consistently the samples when their diversity index values are closer to each other. Still, we found the Chao Shannon entropy to yield the most consistent performances with a Spearman correlation ≥ 0.8 for all tested comparisons. This is depicted Figure 3B, where we averaged the Spearman correlation across each pair of clonal identification methods. It shows that the Chao estimator for Shannon entropy yields the highest averaged correlation over the three datasets (*ρ* = 0.910). Other metrics also reveal high levels of correlation with the exception of the evenness, which only exhibits a correlation of ~ 0.7 (0.756 and 0.617 for the Chao estimator). Evenness being the ratio of two quantities, it is more sensitive to variability in the richness and entropy estimation. Also, the Chao estimator for richness showed lower correlation than the richness itself. As the Chao correction formula relies heavily on the number of identified singletons, a potential cause behind this low performance is the unreliable detection of singletons during the process of clonal identification.

Rather than a single diversity index, the B-cell repertoire landscape may also be characterized in terms of diversity profiles [24] (Figure 3C). Under the Hill’s unified diversity framework [23], the diversity index of order *α* is defined as:

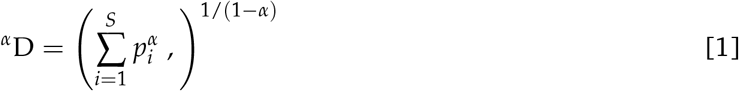

where *p_i_* is the relative abundance of species *i* and ∑ *p_i_* = 1. Values of *α* < 1 tend to favor rare species, while values of *α* > 1 favor the most common species. The interest of Hill’s unified diversity index is that it provides a unified representation of the most common diversity indexes, which can be recovered for different values of *α*, including richness (^0^D), dominance (1/^∞^D), Shannon entropy (log[^1^D]) and the Simpson index (1/^2^D).

We computed the Spearman correlation of the *α* diversity indices across the different clonal identification methods and investigated how the choice of *α* affects the correlation of the diversity indices by setting values of *α* between 0 and 1 with steps of 0.01. We also investigated whether there is optimal *α* that leads to a maximum value of correlation (Figure 3D). Interestingly, the optimal *α* parameter is different for the three datasets studied: *α*_opt_ =0.58 for GC data, *α*_opt_ =0.85 for Hepatitis B, and *α*_opt_ =1.47 for the simulated data. Overall, we found that the value of *α* maximizing the Spearman correlation averaged over the three datasets to be *α*_opt_ =0.97. This is in good agreement with Figure 3B, which indicates that among all the diversity metrics tested, the Shannon entropy and its Chao corrected variant are the optimal indicators. As a reminder, the Shannon entropy *(H)* is closely related to the Hill’s diversity of parameter *α* = 1 (H =log[^1^D]), while other computed indicators are related to ^0^D, ^2^D and ^∞^D.

Finally, we observe a substantial drop in the correlation in Figure 3D for the simulated data. Computing the diversity profile of the simulated repertoires revealed that Hill’s diversity values are near equal across all samples (±1%) for values of *α* ∈ [0.1, 0.9] (Figure S3B). This low variability, possibly coming from an unrealistic simulated environment (not enough variability for the low abundance clones), would explain why the correlation is lower for these values of *α*.

### 1.5 Sensitivity of diversity metrics to sequencing depth

Another question of interest is the sensitivity of these diversity metrics to sequencing depth (or sample size), and whether the traditional Chao statistical estimators are effective in minimizing the bias associated to variability in sample size across samples. To investigate the effect of sequencing depth, we sub-sampled the GC samples with sampling ratios from 1% to 100%. Next, we characterized the clonal populations on each synthetic sample, and evaluated the change in diversity indices across when different clonal identification methods are used. Namely, we computed the fold change between the diversity index values with and without sub-sampling. Figures 4A & B) show the changes in Hill’s diversity index for different levels of sub-sampling. As seen in the figures, changes are consequential for values *α* < 1, which put more weight to rare clonal populations. This finding confirms our expectations, i.e. lower sequencing depths fail to detect rare clonal clusters and result in lower estimations of diversity. On the other hand, no significant change is observed for common clusters, *α* > 1, which are detected even at low sequencing depths. We repeat the analysis using 2 clonal identification methods, the VJ & Junction and the alignment-free method (respectively Figures 4A & B). The same pattern is observed with both methods, with the alignment-free method resulting in a lower range of change between the different sub-sampling ratios. Figure 4C&D show the fold changes between Hill’s diversity index computed with at 10% sub-sampling and 100% sampling, repeated 100 times and averaged over repetitions. *α* = 0 shows the highest variability, which is expected as this metric places equal weight on all clones regardless of their frequency. Singletons, whose detection strongly depends on sequencing depth, contribute to the observed high standard deviation associated with this metric. Similarly to the previous figures, Figure 4C&D indicate that diversity metrics associated with *α* > 1, e.g. dominance, Shannon entropy, Simpson diversity index, etc, are only weakly affected by the sequencing depth.

**Figure 4:**
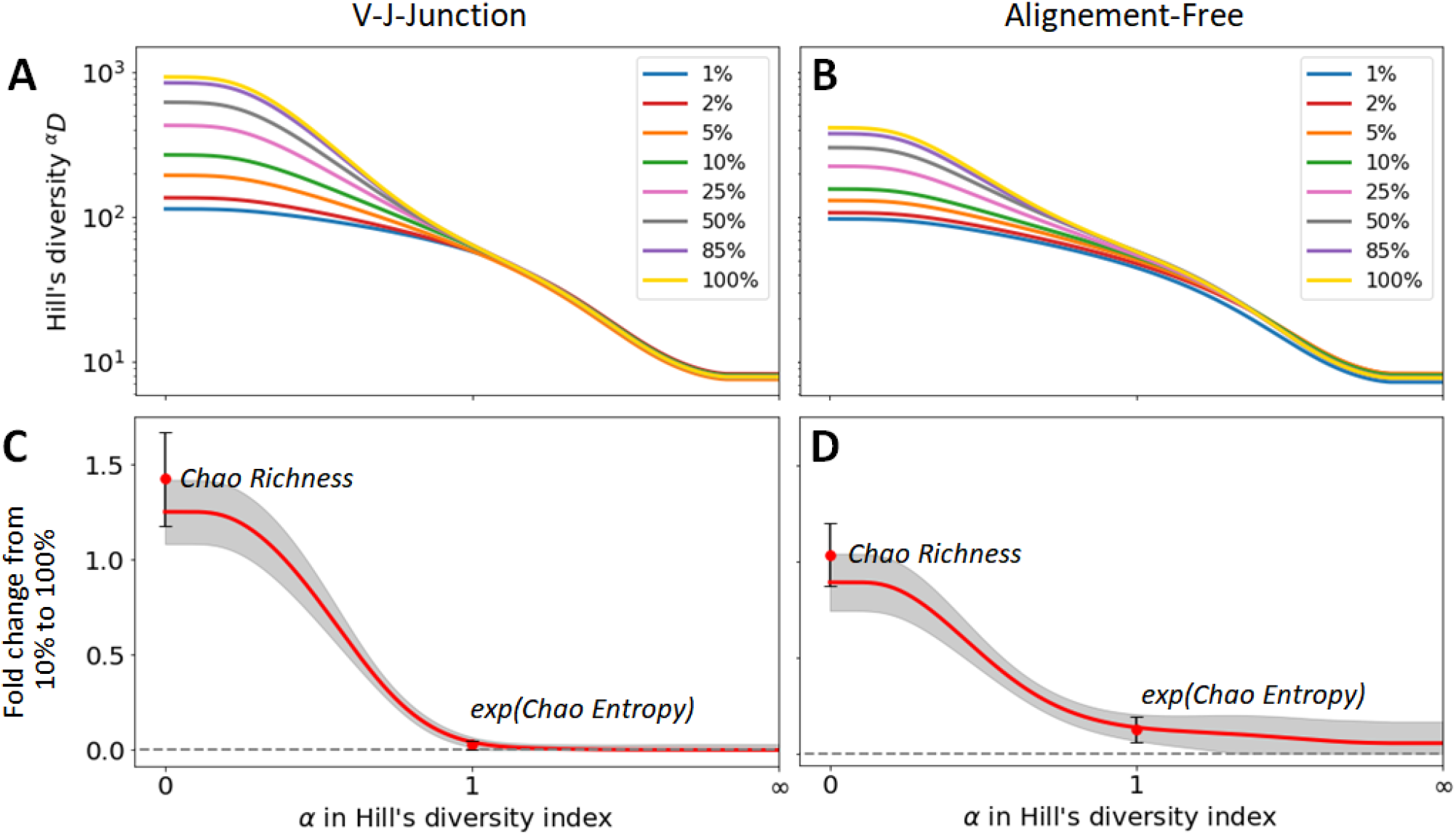
Sensitivity of diversity metrics to sequencing depth for the GC dataset. (A, B) Hill’s diversity profile calculated at different sub-sampling ratios varying from 1% to 100%. (C,D) Fold changes between Hill’s diversity index computed with at 10% sub-sampling and 100% sampling. Mean and one standard deviation across the 100 repetitions of sub-sampling is shown in the figure, with a red line and gray shaded area, respectively. The same results are provided for the richness Chao estimator and entropy Chao estimator (its exponential is shown on for consistency with the Hill’s framework).

The figures also indicate fold changes for the Chao estimators of richness and Shannon entropy (its exponential is shown on Figure 4C&D for consistency with the Hill’s framework, see Method Section 2.4.1). As these indicators are correcting for sample size, we would expect them to be less sensitive to changes in sequencing depth than their uncorrected equivalent (richness and Shannon entropy, respectively). However, this is not what we observe. Interestingly, the Chao estimator for richness even show more sensitivity to sequencing depth (fold change of 1.4) than richness itself (fold change 1.2), while the Chao estimator for Shannon entropy only yielded moderate improvements to the sensitivity to sequencing depth. As the same pattern was observed on the other two datasets, we conclude that the Chao estimators for diversity indexes only result in minor improvements or even make the estimation less accurate. As we discussed in the previous section, this is likely a consequence of unreliable estimation of the number of singletons, which heavily affects the Chao estimators.

## Discussion

B cells play a crucial role in the adaptive immune system, and the advent of efficient high-throughput sequencing of BCRs has generated unprecedented opportunities for the exploration of adaptive immune responses. With these opportunities have come significant challenges in the development of analysis techniques that are robust to technical variations and that can accurately reveal the underlying biological phenomena. The first step in the analysis of B-cell repertoires is the grouping of BCR sequences into B-cell clones that descend from a common ancestor cell. In this study, we compared the performance and potential biases associated with different clone identification methods and highlighted the potential drawbacks of methods relying on germline gene alignments. In fact, we found that these methods may become unreliable when read lengths are shorter than the full V gene (Section 1.3). More importantly, we showed that the choice of the method can greatly impact the inferred clonal structure, especially for low-frequency and singleton clones (Section 1.4). This in turn might bias the analysis of immune repertoires in specific biological contexts.

Our analysis suggests that the VJ & Junction method remains the most accurate to identify clonal groups and singletons, while the junction-only performed worst on simulated data. However, the choice of clonal identification method should be made taking into consideration the experimental design and constraints of each dataset. For instance, we observed that if the V gene assignment is ambiguous, the alignment-free method proposed by Lindenbaum & al. [35] was a better choice to alleviate experimental limitations that result in incorrect V/J assignments. That is because this alignment free method does not rely on the V/J assignments but rather compares the sequence similarity of the whole VDJ sequence by using a vectorized representation of the BCRs. Overall, our results suggest that alignment-free strategies are a promising approach for B-cell clone identification and deserve further investigation.

Another important aspect we explored in this article is the impact of sequencing depth on the quantification of diversity, which is commonly performed to characterize B-cell repertoires. In fact, as the clonal composition across individual’s repertoire is distinct, the translation of these repertoire into several diversity indices offers the unique advantage to extract biological information without directly comparing sequences across repertoires. We performed sub-sampling experiments and characterized the variability of different diversity metrics with sequencing depth and clonal identification methods. We analyzed the change in these metrics when different clonal identification methods were used and found that, while the absolute values were different, different diversity indices including dominance, Shannon entropy, and richness led to similar sample ranks. Shannon entropy was the most robust index (maximizing Spearman sample rank correlation across methods) in the datasets and clonal identification methods we analyzed, which might be due to its weighting similarly both rare and abundant species. Nevertheless, as different diversity indices provide different information, the best practice remains to combine several indices to gain a global view of diversity. In that respect, Hill’s diversity profiles already encompass information about many different indices, and therefore, provides already a more global understanding of B-cell repertoires than any given index. Finally, the use of Chao statistical estimators did not significantly lower the variability of diversity estimation, both in terms of sub-sampling and clonal identification methods. As these estimators rely heavily on singletons and doubletons estimation, a potential cause behind these inefficiencies could be the unreliable detection of singletons during the process of clonal identification.

In summary, we have presented a quantitative comparison of different diversity metrics for the analysis of B-cell repertoires. We have characterized the variability of these metrics when different clonal identification methods were used and for different sequencing depths. One of the main limitations of our analysis is the lack of ground truth in experimental datasets. To partially address this limitation, we have included a synthetic dataset for which the ground truth is known by construction in order to test the different methods. However, addressing this limitation in an experimental context is much harder, and cannot be addressed yet in a fully satisfying manner. Rather, we leveraged *negation sequences* to estimate the specificity of the clone identification methods, i.e. random sequences extracted from different experimental studies, that are very unlikely to be clonally related to sequences in the considered study. However, negation only help in setting the threshold between singletons and non-singletons.

Quantifying the accuracy of identified clones still remains a subjective endeavor. Nevertheless, we grasped an overview of the different methods’ performances by evaluating the agreement between them (AMI). In particular, we checked how singletons predicted by a particular method were treated by the other.

Regarding future work on clonal identification, additional improvements to the method presented in Lindenbaum & al. [35] are promising. For instance, the current alignment-free approach uses BCR vectorized representations based on *k*-mer frequency vectors, as posited by the *tf-idf* metric. This representation does not exploit potential semantic similarities between *k*-mers and assumes that the counts of different *k*-mers provide independent evidence of similarity. Importantly, the order of these *k*-mers in the sequence is not taken into consideration. Furthermore, the frequency vector is dataset dependent, as the *tf-idf* metric computes frequencies across a corpus. Changing the dataset, i.e. the corpus might result in changes in *k*-mers frequencies, and therefore in different clonal groupings. This limits the applicability of the metric across different repertoires. An attractive recent possibility to obtain vectorized representations of BCRs might be to leverage recent neural network and deep learning models for protein tasks, such as Immune2vec [50], ESM [51], TAPE [52], ProGen [53] or ProtBERT [54]. The latent space of these pre-trained models can be used to readily extract a vector representing each BCR sequence. Given the more modest performance of the alignment-free method on the simulated dataset (AMI =0.84) compared to the VJ & Junction method (AMI =0.90), this could be a powerful tool to further improve the accuracy and scalability of current alignment-free clonal identification techniques.

## 2 Methods

### 2.1 B cell repertoire preprocessing

We detail here the preprocessing steps that we performed on all 3 datasets included in our study.

i. Data were downloaded from their original study: GC Data [37] [[link](https://vdjserver.org/community/8899006209436478995-242ac118-0001-012)], hepatitis B vaccination data [6] [[link](https://gateway.ireceptor.org/samples?query_id=67416)], and simulated repertoire data [42] [[link](https://zenodo.org/record/4009638)]. Additionally, a set of the negation sequences was generated by randomly sampling sequences from multiple unrelated individuals [[link](https://vdjserver.org/community/8899006209436478995-242ac118-0001-012)].
ii. For each sequence, the V and J genes were located and annotated based on the alignment to the germline genes downloaded from the IMGT reference directory sets [55], using IgBlast [46].
iii. For the V and J gene assignments, we kept only the germline gene with the highest confidence from IgBlast. In the rare case where a sequence had multiple V and J genes identified with same confidence, we chose the first in alphabetical order.
iv. Sequences were only retained if they were annotated as *productive* by IgBlast.
v. Sequences with the same junction sequences were grouped together and represented by a single sequence randomly selected among them. Sequences within this group were considered to be clonally related because it is very unlikely that two sequences from the different clonal groups have exactly the same junction sequence (SHMs can occur in any part of the sequence and are not limited to the junction region). This assumption greatly reduces the computational workload.

### 2.2 Metrics

#### 2.2.1 Levenshtein distance

The Levenshtein distance [56] is defined as the minimum number of edits required to transform one sequence into another and is a common metric to quantify sequence similarity. To reduce the bias caused by length differences, we used the normalized Levenshtein distance [40] that incorporates the length of both sequences in the following manner:

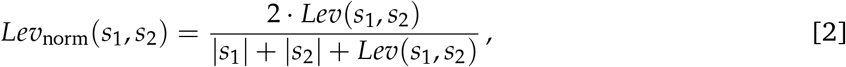

where |*s*_1_| and |*s*_2_| are the lengths of strings *s*_1_ and *s*_2_, and *Lev*(*s*_1_, *s*_2_) is the Levenshtein distance between these two strings.

#### 2.2.2 Cosine similarity

The cosine similarity is a measure of similarity between two vectors. Namely, given vectors **A** and **B**, the cosine similarity is defined as:

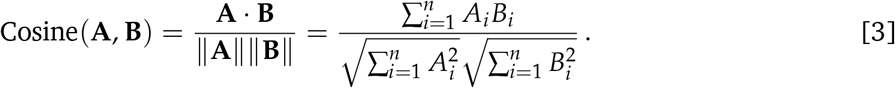

To apply this similarity measure to BCR sequences, we first need to *encode* them. In this paper, we use the term frequency-inverse document frequency (*tf-idf*) weighting scheme.

Briefly, we first compute a *k*-mer representation of each BCR (substrings of length *k*). Then, for each *k*-mer, the frequency term *tf*(*k*) was reweighed with the inverse document frequency which is defined as 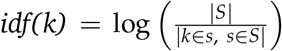, where |*S*| is the total number of sequences and the denominator is the total occurrences of specific *k*-mer *k* across all the *S* sequences. The final *tf-idf* representation was then computed as *tf-idf*(*k*) =*tf*(*k*)·*idf*(*k*). The logic behind this is to emphasize rare and hopefully meaningful terms while reducing the influence of common and uninformative terms.

#### 2.2.3 Adjusted Mutual Information

The mutual information (MI) of two random variables is a measure of the mutual dependence between these two variables. More specifically, it quantifies the “amount of information” obtained about one random variable by observing the other random variable. The mutual information of two jointly discrete random variables X and Y is calculated as:

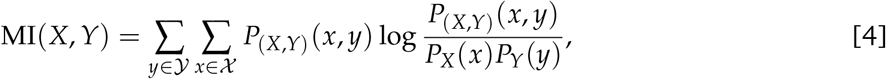

where *P*(*X*,*Y*) is the joint probability mass function of *X* and *Y*, and *P_X_* and *P_Y_* are the marginal probability mass functions of X and Y respectively [57].

MI can also be used to compare clusters, for instance, by measuring the information shared by the two clustering partitions. In practice this is done by counting the number of sequences that are common between each pair of clusters, *A_i_* and *B_j_*, where *A_i_* comes from the first clustering partition 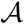 and *B_j_* from the second 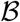:

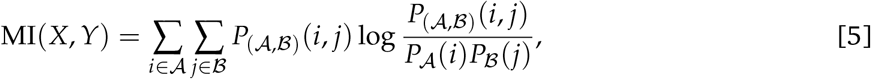

The adjusted mutual information (AMI) is a modified version of the MI to compare two random clusters.

One limitation of the MI to compare partitions is that the baseline value of MI becomes larger when the number of clusters in both partitions increases. To address this limitation, the adjusted mutual information (AMI) can be used instead [44]. Setting *E*{MI(*U*, *V*)} the expected mutual information between two random clusters, the AMI is computed as:

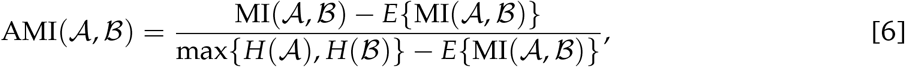

where 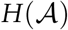 and 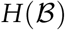 are the entropy associated with the partitioning 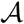 and 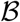, respectively. With this transformation, the AMI takes a value of 1 when the two partitions are identical and 0 when the MI between the two partitions equals the value expected due to chance alone. We used the python implementation from `sklearn` to compute the AMI.

### 2.3 Identifying clones

We implemented three clonal identification methods in this article.

- **Baseline:** B cells were assigned to the same clone if and only if their receptors shared *exactly* the same CDR3 sequence.
- **VJ & Junction:** We first grouped B cells together if and only if they had the same V and J gene. Then for each obtained group, we computed the pairwise normalized Levenshtein distance between each junction in that group, and applied the Hierarchical Agglomerative Clustering (HAC) algorithm [43, 39] to cluster the BCRs into different clonal groups. We used the `complete`-linkage clustering criterion, which begins by clustering each sequence on its own cluster, and then sequentially combines smaller clusters into larger ones until all elements are in the same cluster. The `complete` scheme uses the maximum distances between all observations of the two sets to decide which clusters to merge next. This method results in a dendrogram that shows the sequence of cluster fusion and the distance at which each fusion took place. By setting an appropriate threshold, we can define individual clusters as all the clusters that have not been fusioned up to that distance. In this study, we chose as threshold the distance to the nearest distribution of negation sequences with a tolerance of 1%. Briefly, the threshold is chosen such that it allows a fraction of false-positive sequences roughly equal to a tolerance *δ* to be below the chosen threshold. This heuristic aims for high specificity, which is approximately 1 – *δ*.
- **Alignment-free:** All B cells sequences were first truncated from their *3′* end to a fixed number of nucleotides (*L*), and then encoded into a numerical vector using their k-mer representation reweighted with the term frequency-inverse document frequency (*tf-idf*) weighting scheme (see section 2.2.2). Following on previous work [42], we set *k* = 7 and *L* = 130 as it was found to yield optimal performance in terms of clonal identification. Then, we computed a distance matrix for all sequences in the repertoire using the cosine distance. Cosine distance has the advantage of being very fast to compute for sparse vectors, especially when compared to other alternatives, such as the Euclidean metric. Finally, the threshold definition and clustering into clonal groups were performed using HAC in the same way as in the VJ & Junction method.

### 2.4 Quantifying species diversity

#### 2.4.1 Diversity indexes

Various indexes, such as Shannon entropy [21], Simpson index [22], and species richness [48], are commonly used to quantify the *diversity* of an ecosystem. However, the choice of a universal index to objectively quantify and compare species diversity remains a topic of debate [58]. Starting from the simple assumption that when all species are equally common, diversity should be proportional to the number of species, Hill’s unified diversity framework [23] defines a general formula for the species diversity index that depends on an index *α* as:

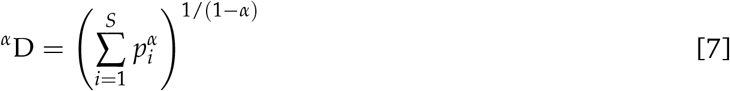

where *p_i_* is the relative abundance of species *i,* and ∑ *p_i_* = 1. For a given number of species *S* > 0, one can prove that 1 ≤ ^*α*^D ≤ *S*.

The choice of *α* plays a role in the weighting of species of different frequencies. *α* < 1 favors rare species, while *α* > 1 favors common species. The most interesting aspect of Hill’s unified diversity index is that one can recover the most common diversity indices used in the literature for particular values of *α*, such as:

- **Species richness** (*α* = 0). The diversity of order zero is insensitive to species abundances and simply corresponds to the number of species: ^0^D =*S*
- **Dominance** (*α* = ∞). Diversity is sometimes represented as the proportion of its most abundant species *i*_max_, corresponding to the inverse of the infinite order diversity index.

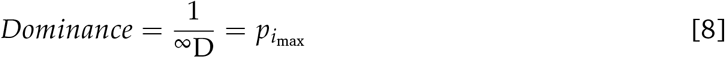
- **Shannon entropy** (*α* = 1).The Shannon entropy (*H*) weighs all species by the log of their frequency. Although Eq.7 is not defined when *α* = 1, its limit exists and converges to the exponential of the Shannon entropy [58].

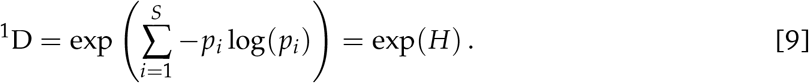
- **Simpson index** (*α* = 2).The Simpson index is defined as:

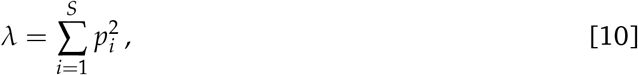

and represents the probability that two entities taken at random from the dataset are of the same type [22]. The Simpson index is directly related to the diversity of order two with 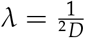.
- **Evenness**. Rather than quantifying the diversity of species, the evenness (E) represents the homogeneity of abundances in a sample or a community [23]. The evenness E(*a, b*) with orders *a* and *b*, *a* > *b*, is defined as

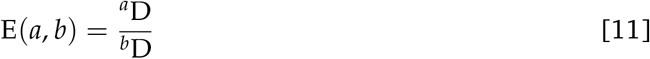 In practice, 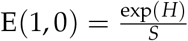 is the most commonly used for quantifying evenness. Note that from this definition we always have 1/*s* ≤ E(*a, b*) ≤ 1. In the case where the number of species is infinite, other values of (*a, b*) should be considered in order to obtain a non-zero evenness [23]. Additionally, the E(1, 0) evenness can be biased when the sample size is small, because it is sensitive to unobserved species. In this case, E(2, 1) is preferred.

#### 2.4.2 Hill’s Diversity profile

Because Hill’s diversity profiles encompass the information contained in several diversity indices, its use is becoming increasingly common to obtain fingerprints of the immune repertoire [23, 24].

In this paper, we treated the *diversity profiles* as an *N* dimensional vector, where each element of the vector is the Hill’s diversity index ^*α*^D for a different value of *α* ∈ [0, ∞]. Starting from a vector **A** with *N* elements in the range [−1, 1], we obtain the values of ***α*** = [*α*_1_ ··· *α_N_*] for our diversity profile with the transformation.

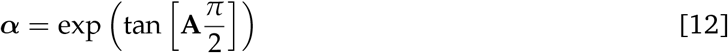

Then, the diversity profile ^*α*^D is computed as

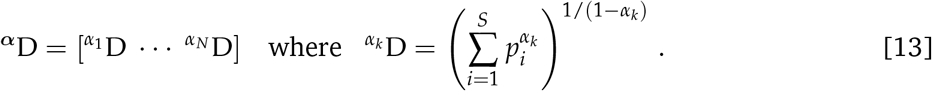

We introduced that transformation to (i) be able to include both the Richness (^0^D) and Dominance (1/^∞^D) in a finite vector, and (ii) to respect the symmetry between in the weighting of species of different frequencies (where *α* < 1 favors rare species, while *α* > 1 favors common species).

#### 2.4.3 Estimating diversity with incomplete sample information

Complete knowledge about a system is often not available. Partial knowledge often results in the underestimation of a sample’s diversity, as some species might not have been observed. Specialised statistical tools have been developed to estimate the true richness *S*_true_, i.e. the true number of species, of a sample. One of the most common is the bias-corrected Chao1 species richness estimator [25, 26].

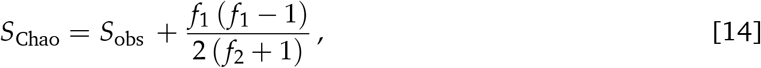

where *S*_obs_ is the total number of species detected, *f*_1_ is the number of species detected exactly once, and *f*_2_ the number of species detected exactly twice. The intuition behind this indicator is that if many species are detected only once, there is likely a large number of species that have not yet been detected. On the other hand, when all species have been detected at least twice, it is unlikely that new undetected species exist.

In addition to the Chao1 estimator for species richness, a similar approach can be used to estimate the Shannon entropy with incomplete sample information [49]. Defining *n* as the number of observations, we can estimate the sample coverage as 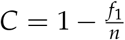 - this equation represents a first order approximation based only on singletons, and adjust the relative species abundance with 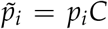. Then the Chao estimator for Shannon entropy can be defined as:

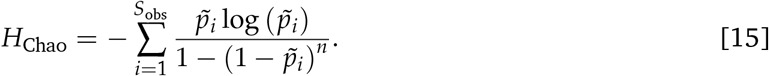

Estimators can also be computed for higher orders of diversity, for instance, using the general Horvitz-Thompson estimator [59] or other Chaos estimators such as diversity rarefaction curves [27].

## Author Contributions

S.L. wrote the code and analyzed the data under the supervision of A.P. and M.R.M. S.L. and A.P. wrote the manuscript. All authors have read and agreed to the published version of the manuscript.

## Code and Data Availability

All the code used in our study is available at https://github.com/Aurelien-Pelissier/cdiversity. We make our implementation freely available as a Python library -`pip install cdiversity`. The dataset used in this study are available at:

- Germinal center data [37] [[link](https://vdjserver.org/community/8899006209436478995-242ac118-0001-012)]
- Hepatitis B vaccination data [6] [[link](https://gateway.ireceptor.org/samples?query_id=67416)]
- Simulated repertoire data [42] [[link](https://zenodo.org/record/4009638)].

