## Supplementary Materials for "Quantifying B-cell Clonal Diversity In Repertoire Data"

[*] *Equal contribution*

### 1 Distance to nearest distribution for each dataset

The distance to nearest distribution is a key component in the identification of clones, as it is required to define the right threshold for the clustering. Unsurprisingly, the shape of the distribution, as well as the bi-modality of singletons and non-singletons looks different depending on the metric used (Figure 1).

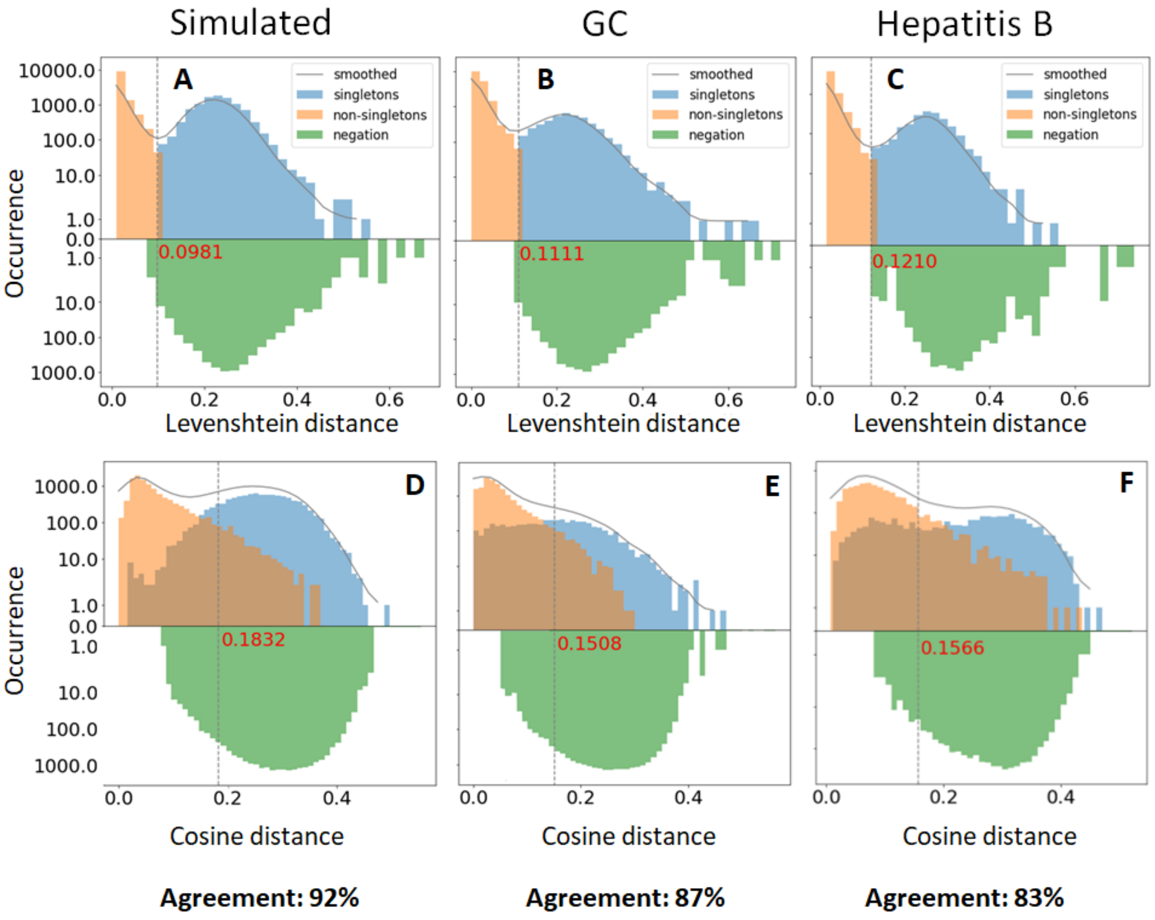


**Figure 1:** Distance to nearest distribution for each dataset in the context of both the alignment-based (Levenshtein distance) and alignment-free (Cosine distance) clonal identification method. In order to visualize the *agreement* between the two method (percentage of equal prediction), the singletons predicted by the first methods are used to label the distributions on the second method. The distance to nearest sequence in the negation dataset is also shown as a reference point. It was set with a tolerance of 1% for the alignment-free method

Interestingly, a subset of predicted singletons with a low Levenshtein distance in the alignment-based method have a high distance in the context of the alignment-free method. As a result, the singletons identified by both methods do not match exactly (percentage of equal prediction of 92%, 87% and 83% respectively across each dataset).

### 2 Robustness analysis of clonal identification methods

One question of interrest is how uncertain the clone assignment of a BCR sequence is in terms of the randomness in biological sampling. Since the comparison across different cell populations is non-trivial, we investigated the consistency of the clone identifier by comparing the clustering of the same subset of sequences, but with/without the accompanying of the rest sequences as input. To do so, we sub-sampled each sample by randomly selecting half of its sequence, and then performed the clustering again with the subsampled sequences only (Figure 3A). After that, we compared the obtained clustering results to the original one (with the full sample) with the adjusted mutual information (AMI). In this case, the computed AMI represents the *robustness* of the clonal identification methods to biological sampling.

We performed this analysis for each sample and depicts the results Figure 3B. On average, the AMI of the junction-only, alignment-based and alignment-free method were $1, 0.92$, and $0.88$ respectively. While our analysis indicates that all three methods have highly consistent clonal identification results, there is still significant differences between the three methods. As expected, the junction-only method always yields exactly the same results, since it do not rely on any clustering techniques. On the other hand, the alignment-free method is the least robust, as the BCR vectorized representation of sequences themselves changes with the subsampling (due to the learned *idf* reweighting).

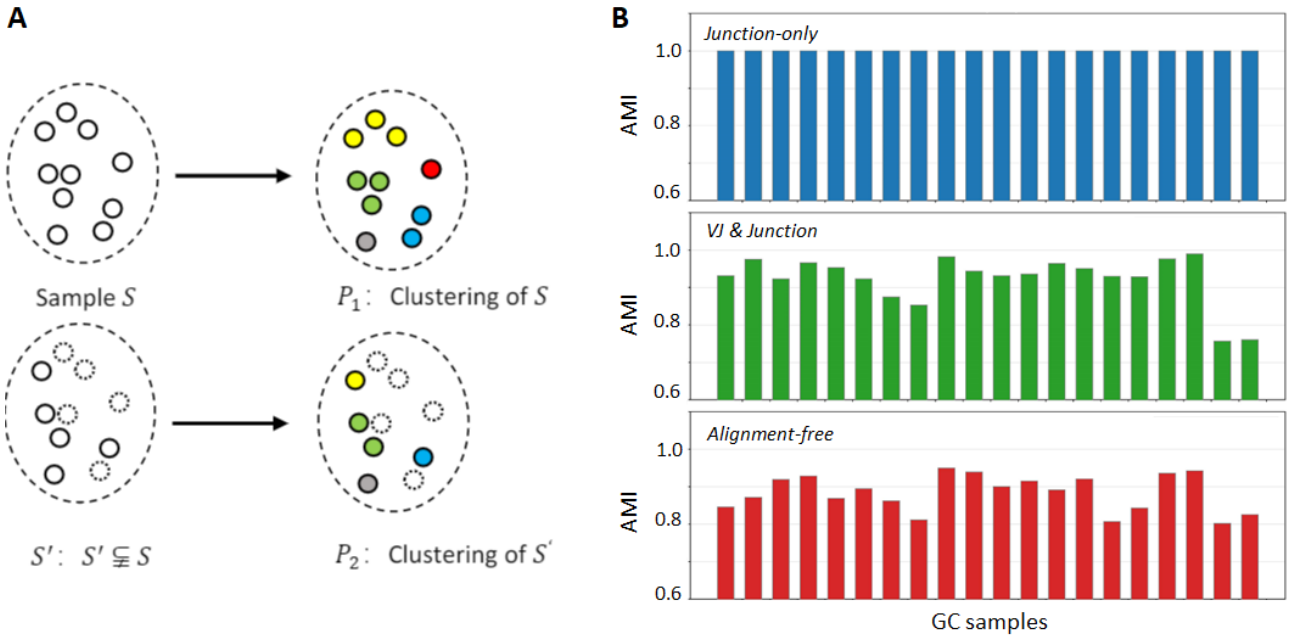


**Figure 2:** Robustness analysis of clonal identification methods. (A) A subsample $S'$ is generated from sample $S$, and the obtained clonal families are compared across the two clustering. (B) Adjusted mutual information (AMI) between the original and subsampled clustering across each sample in the GC dataset, for the three clonal identification methods.

#### 3 False negative singletons with the Alignment-free method

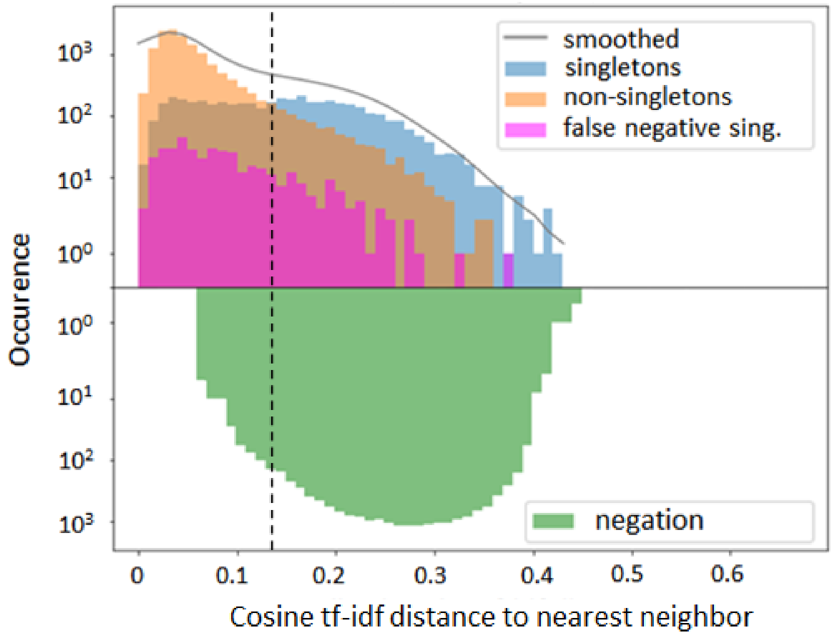


**Figure 3:** Distance to nearest distribution for B-cell sequences in the alignment free clonal identification method in the GC data. The sequences (singletons and non-singletons) are labeled with the alignment-based (VJ & Junction) method, where the false negative singletons were found from the ambiguous V or J gene misalignments (depicted in purple).

#### 4 Spearman correlation of diversity indices across methods

In Table 1, we provide the Spearman rank correlation for diversity indices in different biological context. First, we compare the diversity indices across the different clonal identification methods, which can be performed on the three datasets and that we extensively discuss in the main text. In the specific case of the simulated data, we can also test for the correlation between the diversity indices from different clonal identification methods and to ground truth diversity. Here we see that Shannon Entropy seems to perform the best in the context of different clocal identification. Finally, in the specific case of the germinal center dataset, we can test for the consistency of the diversity indices across replicates for each clonal identification method (we have two replicates per GC). The results Table 1 suggest that the Simpson index seems to be optimal in the context for the GC replicates (maximizing correlation averaged over the three replicates).

| | Diversity index | JO *vs* VJJ | JO *vs* AF | VJJ *vs* AF | JO *vs* $G_0$ | VJJ *vs* $G_0$ | AF *vs* $G_0$ | *Mean* |
|---|---|---|---|---|---|---|---|---|
| Simulated dataset | Richness | 0.89 | 0.82 | 0.84 | 0.79 | 0.82 | 0.84 | *0.83* |
| | Chao Richness | 0.69 | 0.64 | 0.56 | 0.63 | 0.89 | 0.72 | *0.69* |
| | Shannon Entropy | 0.92 | **0.88** | 0.70 | 0.69 | 0.88 | 0.84 | *0.82* |
| | Chao Shannon Entropy | **0.96** | 0.80 | 0.83 | 0.84 | **0.91** | **0.89** | *0.87* |
| | Simpson index | 0.94 | 0.70 | 0.69 | 0.65 | 0.75 | 0.64 | *0.73* |
| | Dominance | 0.82 | 0.87 | 0.70 | 0.72 | 0.79 | 0.80 | *0.78* |
| | Evenness | **0.96** | **0.88** | **0.84** | **0.85** | **0.91** | 0.87 | ***0.89*** |
| | *Mean* | *0.88* | *0.80* | *0.74* | *0.74* | *0.85* | *0.80* | |

| | Diversity index | JO *vs* VJJ | JO *vs* AF | VJJ *vs* AF | repl. JO | repl. VJJ | repl. AF | *Mean* |
|---|---|---|---|---|---|---|---|---|
| GC dataset | Richness | 0.79 | **0.97** | 0.76 | 0.47 | 0.94 | 0.90 | *0.81* |
| | Chao Richness | 0.82 | 0.96 | 0.77 | 0.56 | 0.64 | 0.81 | *0.76* |
| | Shannon Entropy | **0.99** | 0.91 | **0.89** | 0.85 | **0.96** | 0.82 | *0.90* |
| | Chao Shannon Entropy | **0.99** | 0.90 | **0.89** | 0.85 | **0.96** | 0.84 | *0.91* |
| | Simpson index | 0.97 | 0.90 | 0.88 | **0.95** | 0.92 | **0.96** | ***0.93*** |
| | Dominance | 0.91 | 0.92 | 0.86 | 0.88 | 0.93 | 0.95 | *0.91* |
| | Evenness | 0.63 | 0.71 | 0.52 | 0.66 | 0.75 | 0.61 | *0.65* |
| | *Mean* | *0.87* | *0.90* | *0.80* | *0.75* | *0.87* | *0.84* | |

| | Diversity index | JO *vs* VJJ | JO *vs* AF | VJJ *vs* AF | *Mean* |
|---|---|---|---|---|---|
| Hepatitis B dataset | Richness | 0.98 | 0.98 | 0.94 | *0.97* |
| | Chao Richness | 0.87 | 0.96 | 0.80 | *0.88* |
| | Shannon Entropy | **1.00** | **0.99** | **0.99** | ***0.99*** |
| | Chao Shannon Entropy | **1.00** | **0.99** | **0.99** | ***0.99*** |
| | Simpson index | **1.00** | **0.99** | **0.99** | ***0.99*** |
| | Dominance | **1.00** | 0.98 | 0.97 | *0.98* |
| | Evenness | 0.99 | **0.99** | 0.96 | *0.98* |
| | *Mean* | *0.98* | *0.98* | *0.95* | |

**Table 1:** Spearman correlation of the diversity indices values across samples for each dataset. The correlation is computed between each pair of clonal identification method. *Junction-only* (JO), *VJ & Junction* (VJJ) and *Alignment-free* (AF). For the simulated dataset, comparisons are also performed on ground truth ($G_0$) clonal groups. For the GC dataset, we also compute the correlation between the GC replicates (designated as repl). Max values for each columns are highlighted in bold

#### 5 Diversity profiles of each dataset

We computed the Shannon entropy of each sample in each dataset and we show the results Figure 4A. Clearly, the variability of the Shannon diversity relative to the average value is not the same across sample. It seems that the simulated dataset show small variability in the Shannon entropy (mean/std $\sim 80$) while the Hepatitis B and GC dataset show higher variability (mean/std $\sim 3$ and $\sim 1.5$, respectively). This is reflected on Figure 4B, where the Hill's diversity profiles becomes near superposed for $\alpha < 1$. In the context of low variability, ranking the samples Hill's diversities for these values of $\alpha$ might not be consistent.

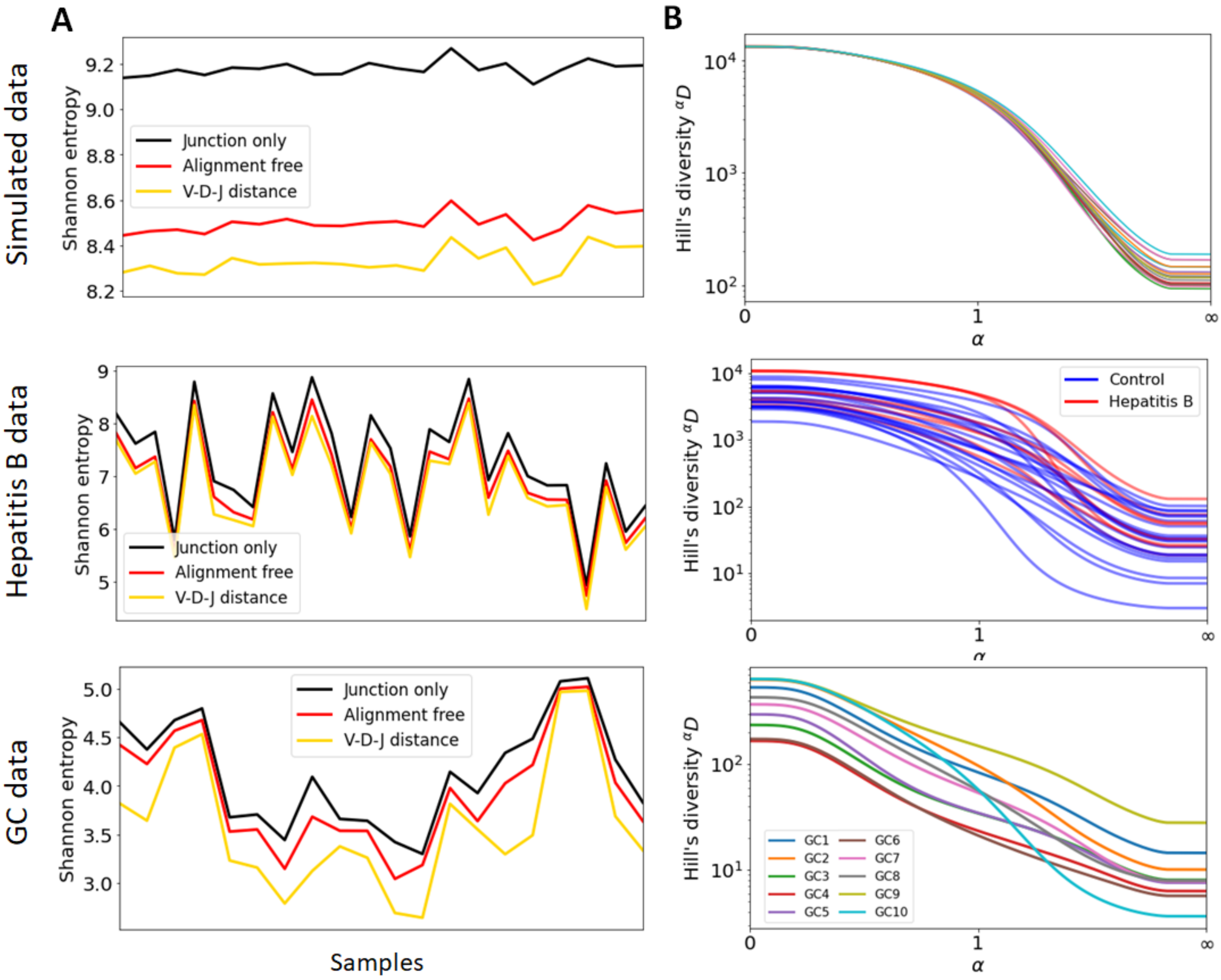


**Figure 4:** B-cell repertoire clonal diversity analysis. (A) Shannon entropy of each sample in each dataset for the three clonal identification methods. (B) Hill's diversity profile of each sample in each dataset, with clones inferred from the alignement-free method. Note that the $x$ axis was transformed by a exponential tangent function for visual clarity.
